# Homemade Bread: Repurposing an Ancient Technology for *in vitro* Tissue Engineering

**DOI:** 10.1101/2020.11.13.353698

**Authors:** Jessica T. Holmes, Ziba Jaberansari, William Collins, Maxime Leblanc Latour, Daniel J. Modulevsky, Andrew E. Pelling

## Abstract

Numerous synthetic and naturally-occurring biomaterials have been developed to provide such architectures to support the proliferation of mammalian cells in vitro and in vivo. Our group, and others, have shown that scaffolds derived from plants can be utilized for tissue engineering applications in biomedicine and in the burgeoning cultured meat industry. Such scaffolds are ideally straightforward and inexpensive to prepare, allowing researchers to take advantage of their intrinsic 3D microarchitectures. These efforts inspired us to continue to pursue the development of novel and unconventional biomaterials that are easily produced and high performing *in vitro*. With this in mind, few plant-derived materials are more ubiquitous than bread. Having observed the porosity of the crumb (i.e. the internal bulk) we sought to investigate whether it might support the proliferation of mammalian cells *in vitro*. Here, we develop and validate a yeast-free “soda bread” that maintains its mechanical stability over several weeks in culture conditions. Importantly, we also demonstrate that control over the mechanical stability of the scaffold can also be achieved with both chemical and enzymatic means. The scaffolding is a heterogeneous and complex structure of isolated and interconnected pores which allow for the proliferation of multiple cell types. We demonstrate here that mouse fibroblasts, myoblasts and pre-osteoblasts are able to proliferate up to four weeks in culture. Immunohistochemistry demonstrates that the fibroblasts are able to deposit their own fibronectin extracellular matrix and that mouse myoblasts are able to differentiate and fuse into myotubes. Although the pre-osteoblasts proliferated over the course of four weeks their ability to differentiate was inconclusive. Metabolic analyses of proliferation, cytotoxicity and oxidative stress reveal that cells remain highly viable and functional on these novel bread scaffolds. While the results presented in this proof-of-concept study create many new questions and opportunities, the results open up novel possibilities in the development of edible scaffolds that may be utilized in future food applications. Bread derived scaffolds represent a surprising alternative to synthetic or animal-derived scaffolds for addressing a diverse variety of tissue engineering challenges in food science. Future studies will delve deeper into investigating these how possibilities might take advantage of the immense breadth of knowledge about bread making and examine their applicability in the development of lab grown foods and broader applications in cellular agriculture.

## 1. INTRODUCTION

In recent years, there has been an increase in studies on the use of plant-derived biomaterials for tissue engineering applications [1–12]. For example, in work from our group we were able to demonstrate that decellularization of plant tissues resulted in cellulose-rich three dimensional (3D) scaffolds [1–3]. In addition, we have demonstrated that these scaffolds also perform very well after implantation into animal models, resulting in a high degree of tissue integration and vascularization [2,3]. Several other groups have also published similar studies with a multitude of plant tissues and mammalian cell types to demonstrate the general utility of plant-derived biomaterials for biomedical and food-based tissue engineering applications [5–12]. A growing body of literature also suggests that plant-derived proteins can be utilized to create scaffolds for tissue engineering and are broadly compatible with mammalian cell culture. Proteins such as soy, zein and camelina, etc have been studied, but of particular interest are gluten proteins derived from wheat, such as gliadin and glutenin [13–15]. Such wheat derived proteins can be purified and made into films suitable to culture mammalian cells. For instance, glutenin films have been demonstrated to be an acceptable substrate for osteoblasts [14]. In the same study, a gluten film was shown to support the growth of osteoblasts but with less efficiency due to the cytotoxicity of gliadin [14]. Wheat protein based scaffolds can also be obtained through electrospinning, in which ultrafine fibrous structures can be obtained, creating a polymer melt film of wheat glutenin [15]. Such scaffolds have been shown to support the culture of adipose derived mesenchymal stem cells [14]. Although effective, these methods are labor and resource intensive, requiring two days to purify the proteins and seven days for them to be electrospun [14,15].

All things considered, the development of naturally derived 3D biomaterials has gained considerable interest in recent years due to their potential for use in cellular agriculture applications [5,12,16]. Although still emerging, the broader goal of cellular agriculture [17–19] is to replace products produced by traditional agricultural methods with biotechnological approaches notably through synthetic biology and tissue engineering [17,18]. One specific focal point within the field is the cultivation of mammalian cells *in vitro* for the preparation of meat-like products [5,17,20]. Although a number of challenges remain to achieve this goal, a significant body of work has begun to address issues such as the large scale production of relevant cell types, creating sustainable and ethically sourced media and developing suitable scaffolds [5,17,18,20]. While exciting, it is important to also recognize that the future potential of these methods are still debated as efforts to address the continued dependence on animal products in traditional cell culture (for example, fetal bovine serum), water/electricity use and any potential health benefits in the final meat products are still ongoing [21]. With that said, there remains an intense interest in developing solutions which address these issues, and others, potentially opening up a new future in which foods can be more sustainably created and distributed globally. Plant-derived scaffolds are of particular interest due to the potential of creating edible scaffolds [5,12,16]. Inspired by the simplicity of plant-based biomaterials, we investigated the possibility of developing another complementary approach to fabricating scaffolds for tissue culture which is simple and straightforward to implement.

The work presented here was strongly influenced and motivated by recent global events, namely the SARS-CoV-2 pandemic. As many people around the world found themselves physically distancing and working from home, baking bread rapidly gained a large increase in popularity. According to Google Trends, the search queries for the term ‘bread’ spiked in the early days of the global pandemic [22]. As with many other laboratories around the world, our laboratory was shut down during this same period. As a research group, many of us found that we were also participating in our own efforts to bake bread at home. Sharing our experiences, we hypothesized that baked bread might also possess the structural characteristics to make it suitable as a 3D, porous and biocompatible biomaterial for mammalian cell culture applications. During that time we designed a series of experiments which have since been performed, the results of which we present below. The main ingredient, wheat flour, is primarily composed of starch (~70%) but its strength and stability is mainly attributed to its protein content (~12%), represented by gluten and non-gluten (albumin and globulins) proteins [23]. As described above, such proteins have found potential utility in the creation of biomaterials [13,15].

A requirement for an optimal biomaterial is a structure with high porosity to prevent an anoxic microenvironment. The porous nature of bread results from the presence of a leavening agent, such as yeast or baking powder. The reaction between sodium bicarbonate, the active ingredient in baking powder, and water yields bicarbonate, an anion that decomposes into water and carbon dioxide at ambient temperature and is further favoured when exposed to heat. The carbon dioxide helps the bread rise in addition to leaving behind pores as it exits the crumb. The porous structure of the crumb allows for the migration of cells within the bread which consequently makes it an appealing biomaterial. Moreover, the time required to make the bread is less than an hour, which is highly efficient in comparison to other methods of creating scaffolds for tissue engineering, including plant-based scaffolds. The ingredients needed to make the dough are found in most household pantries and can be purchased at a fraction of the cost of the chemicals often required to fabricate other types of biomaterials.

In this study, we present a soda bread recipe that can be used as an easily produced scaffold that supports the proliferation of mammalian cells in culture. Our objective was to demonstrate that mammalian cells could remain viable and proliferate *in vitro* within scaffolds made of porous bread crumb. We demonstrate that bread scaffolds remain intact over the course of up to four weeks in cell culture, can be modified to control their mechanical properties and that mammalian cells will proliferate and remain viable within the scaffolds. Importantly, the recipe we utilize relies on sodium bicarbonate, rather than yeast, to create the required porosity, thus avoiding potential contamination of the scaffold with an unwanted cell type. The data presented here supports a simple and highly accessible new method for creating cell culture scaffolding utilizing ancient approaches. We demonstrate below that several cell types can be cultured on these scaffolds and that they remain viable and do not exhibit signs of significant oxidative stress or cytotoxicity. Furthermore, we demonstrate that mouse myoblasts are also capable of differentiating and fusing into multinucleated myotubes on the scaffolds. The inherent porosity of the bread crumb along with its simple production method results in a potentially useful biomaterial that can be utilized for *in vitro* 3D cell culture. Time will be required to fully elucidate how such a scaffold may be effectively scaled in an industrial cultured meat setting; however, the results uncover a novel potential path forward towards the goal of meat production in an animal free manner. Future work will build upon this first proof-of-concept which established the use of bread as scaffolding biomaterial to further present methods for cell type specific optimization of surface chemistry, physical properties, and porosity/structural control. Regardless, the research presented here establishes a novel approach for developing edible biomaterials that can potentially be employed in the production of laboratory grown meat and other cellular agriculture applications.

## 2. MATERIALS AND METHODS

### 2.1 Bread Recipe and Fabrication

Our approach is based on a common soda bread recipe. In a ceramic bowl, 120g of all purpose flour (Five Roses), 2g of iodized table salt (Windsor) and 10g of baking powder (Kraft) were mixed together. This was followed by the addition of 70 mL of water, which was heated for 30 seconds in a microwave until the temperature of the water was ~75°C. The mixture was combined to form a dough and shaped into a ball. The dough is then kneaded for 3 minutes with the addition of flour as needed to reduce sticking. Once flattened into a circular disk with a height of approximately 2.5 cm, the dough was placed in a glass bread pan lined with parchment paper. It was baked for 30 minutes at 205°C in a preheated oven. The cooled bread was stored in a resealable plastic bag (Ziploc) at −20°C until use.

When ready for use, the bread was thawed to room temperature. A 6mm biopsy punch is then be used to extract cylindrical samples from the internal portion of the loaf (ie. the “crumb”). The cylinders are then cut with a blade (Leica) to form circular scaffolds, which are about 2.5mm in thickness. In some cases, scaffolds were crosslinked with glutaraldehyde (GA) to stabilize their mechanical properties. In this case, we adapted an approach described previously for similar protein based scaffolds [24,25]. A 0.5% GA solution was prepared from a 50% electron microscopy grade glutaraldehyde stock (Sigma) and diluted with PBS (Fisher). The scaffolds were incubated in the GA solution overnight in the fridge. Afterwards, the scaffolds were rinsed 3 times with PBS. To reduce any remaining traces of unreacted glutaraldehyde, the scaffolds were incubated in a 1 mg/mL NaBH4 (Acros Organics) solution on ice, made immediately before use. Once the formation of bubbles ceased, the samples were rinsed 3 times with PBS. In some cases, the bread scaffolds were also crosslinked with transglutaminase (Modernist Pantry). TG is a well known enzyme that catalyzes protein crosslinking by forming covalent links between the carboxamide and amino groups of glycine and lysine respectively. Here, a pilot study was performed in which TG was mixed with the dry ingredients at a concentration of 1% (w/w) in advance of baking.

### 2.2 Cell Culture

NIH3T3 mouse fibroblasts stably expressing GFP were used in this study (ATCC). Cells were cultured in high glucose Dulbecco’s Modified Eagle Medium (DMEM) (HyClone), supplemented with 10% fetal bovine serum (HyClone) and 1% penicillin/streptomycin (HyClone) at 37°C and 5% CO_2_. The culture media was exchanged every second day and the cells were passaged at 70 % confluence. To test the suitability of the scaffold to support the proliferation of other cell types, C2C12 mouse myoblasts and MC-3T3 mouse pre-osteoblasts were also cultured on the scaffolds according to the protocols above. In the case of MC-3T3 cells, the DMEM was replaced with Minimum Essential Medium (MEM) (ThermoFisher).

To prepare bread scaffolds for seeding, they were placed in 70% ethanol for 30 minutes in order to sterilize them and then rinsed twice with PBS. Bread scaffolds were additionally soaked in complete media prior to seeding to encourage adherence. A droplet containing 1.0×10^5^ cells was then gently placed on top of each scaffold which were contained in 12-well plates. The samples were placed in the incubator for 3-4 hours to allow the cells to adhere to the scaffolds. Afterwards, 1.5-2 mL of culture media was added to each well and the samples were incubated. The culture media was exchanged every 48-72 hours. Cells were maintained on scaffolds for two weeks in a standard cell culture incubator.

Differentiation of C2C12 and MC-3T3 cells was also carried out as part of this study. C2C12 differentiation was initiated after first allowing the cells to grow to confluence over a period of two weeks. At this point cells were cultured in myogenic differentiation media (DMEM, 2% Horse Serum, 1% penicillin/streptomycin) for up to two weeks in order to stimulate cell fusion and myogenesis. MC-3T3 cells were differentiated following a similar protocol but with osteogenic differentiation media (MEM, 10% fetal bovine serum, 1% penicillin/streptomycin, 50 μg/mL ascorbic acid and 10 mM β-glycerophosphate) for up to four weeks.

### 2.3 Staining

Before staining, the scaffolds were fixed in 4% paraformaldehyde for 10-15 minutes at room temperature. Following 3 rinses with a duration of 5 min each in PBS, samples containing NIH3T3-GFP cells were stained using a stock Hoechst 33342 (ThermoFisher) stock solution (1:500 in PBS) for 15 minutes to label nuclei. In cases where C2C12 and MC-3T3 cells were cultured, after fixation with 3.5% paraformaldehyde the cells were permeabilized with Triton X-100. Phalloidin alexa fluor 488 (ThermoFisher) stock solution (1:100 in PBS) was incubated on the samples for 20 min at room temperature to label actin. In cases of antibody staining, samples were first washed with an ice cold wash buffer (PBS, 5% FBS, 0.05% sodium azide) and placed on ice. C2C12 myotubes were labeled by incubating with an MF-20 myosin heavy chain primary antibody at a 1:200 dilution (DSHB Hybridoma Product) for 30 min followed by a rat anti-mouse IgG secondary antibody conjugated to Alexa Fluor 488 at a 1:100 dilution for 30 min. Between each stain the sample was incubated with the wash buffer for 30 min and the entire process was carried out on ice. In cases where deposited fibronectin was labelled the process was similar to the above. However, samples were incubated with a primary anti-fibronectin antibody at a 1:200 dilution (Abcam) for 30 min, followed by a rabbit anti-mouse IgG secondary antibody conjugated to Alexa Fluor 546 at a 1:100 dilution for 30 min. After any staining protocol, all scaffolds were rinsed for 2 min with PBS. The scaffolds themselves were then stained with either a 0.2% (m/m) Congo Red (Sigma) stock solution, or a 50% (v/v) dilution of Calcofluor White stock solution (Sigma), for 15 minutes. This was followed by 5-10 minute washes with PBS prior to mounting and imaging.

### 2.4 Confocal Microscopy

Confocal images were obtained using an A1R high speed laser scanning confocal system on a TiE inverted optical microscope platform (Nikon, Canada) with appropriate laser lines and filter sets. Images were analyzed using ImageJ open access software (http://rsbweb.nih.gov/ij/). Brightness and contrast adjustments were the only manipulations performed to images.The ImageJ software was also used to count the number of cells in different areas of the scaffolds. Image analysis was conducted for quantifying pore size and volume fraction by collecting confocal Z-stacks, applying a threshold to obtain binary images at each optical plane, denoising and image quantification of pore area and volume.

### 2.5 Scanning Electron Microscopy

The preparation of the samples for SEM began with a fixation in paraformaldehyde. This was followed by a dehydration through successive washes of ethanol with increasing concentration (35%, 50%, 70%, 95% and 99%). The samples are dried using a PVT 3D critical point dryer (Tousimis) and gold-coated at a current of 15 mA for 3 minutes with a Hitachi E-1010 ion sputter device. SEM images were acquired at a voltage of 2.00 kV on a JEOL JSM-7500F FESEM. In the case of scaffolds cultured with MC-3T3 cells, energy-dispersive spectroscopy (EDS) was performed on three different areas of each scaffold surface and analyzed for mineral aggregates.

### 2.6 Cell Viability Assay

Cell viability was assessed with the Alamar Blue assay (Invitrogen). Cells were seeded onto scaffolds and assessed after 1 and 13 days in culture. In each case, samples were incubated with 10% (v/v) Alamar blue solution standard culture media for 2h in the incubator. Following incubation, fluorescence was measured in a microplate reader at 570nm against reference wavelength at 600nm. The results are expressed in arbitrary units (AU) and normalized against the initial readings after 1 day in culture of the respective cell types.

### 2.7 Glutathione Assay

A Glutathione Assay (Cayman Chem) was conducted to evaluate the abundance of antioxidants within cells following incubation. Following two weeks of incubation the NIH3T3, C2C12 and MC-3T3 cultures were evaluated for glutathione content according to the manufacturers guidelines. In brief, both 2D cell culture and 3D bread samples were collected and lysed in 2-(N-morpholino)ethanesulfonic acid (MES) buffer and centrifuged at 10,000 × g for 15 minutes and then deproteinated with metaphosphoric acid (MPA). The resulting lysates were quantified against a standard curve as described by the supplier and normalized against the protein content of each sample by bradford assay.

### 2.8 Lactate Dehydrogenase Cytotoxicity Assay

Cytotoxicity was evaluated using the CyQuant LDH Cytotoxicity Assay (ThermoFisher) to evaluate cell health. Samples of NIH3T3, C2C12 and MC-3T3 cells were incubated for two weeks in culture as described previously and compared against 3D TCP controls. Both ‘Spontaneous’, what is released in culture, and ‘Maximum’, the maximal value following lysis, fractions were collected to provide a %-Cytotoxicity value as described by the manufacturer. Results are expressed as the difference between the spontaneous and max reported values comparing the 2D TCP and 3D bread experimental conditions.

### 2.9 Mechanical Testing

The Young’s modulus of the scaffolds was determined by compressing the material up to a maximum of 85% strain, at a rate of 3 mm/min, using a custom-built mechanical tester controlled with LabVIEW software. The force–compression curves were converted to stress–strain curves and the slope in a linear regime was fit (typically between 10-20% compression) to extract the Young’s modulus.

### 3.0 Statistics

For the comparison of time series data, a one way ANOVA with Tukey’s post-hoc analysis was used to determine the statistical difference between sample populations. To compare between two distinct populations a student’s t-test was employed. In all cases alpha = 0.05. Where indicated, all values are presented as the mean ± standard deviation. Analysis and statistical tests were conducted using the OriginLab software package.

## 3. RESULTS

### 3.1 Preparing sterile bread-derived scaffolds

Scaffolds were fabricated as described in the methods section and in a manner similar to many bread recipes. First, dry ingredients were combined followed by mixing in warm water and kneading (Fig. 1A, D). After baking the internal part (crumb) of each loaf was characterized by a network of material which possessed a significant variability in its porosity (Fig. 1C, D). To prepare the bread as a scaffold for supporting cell culture, a 6mm biopsy punch was used to extract a cylinder of material from the internal portion of the loaf, also known as the crumb. The cylinder was then sliced with a scalpel to create approximately 2.5mm thick, 6mm diameter circular pieces of material (Fig. 1D).

**Figure 1.**
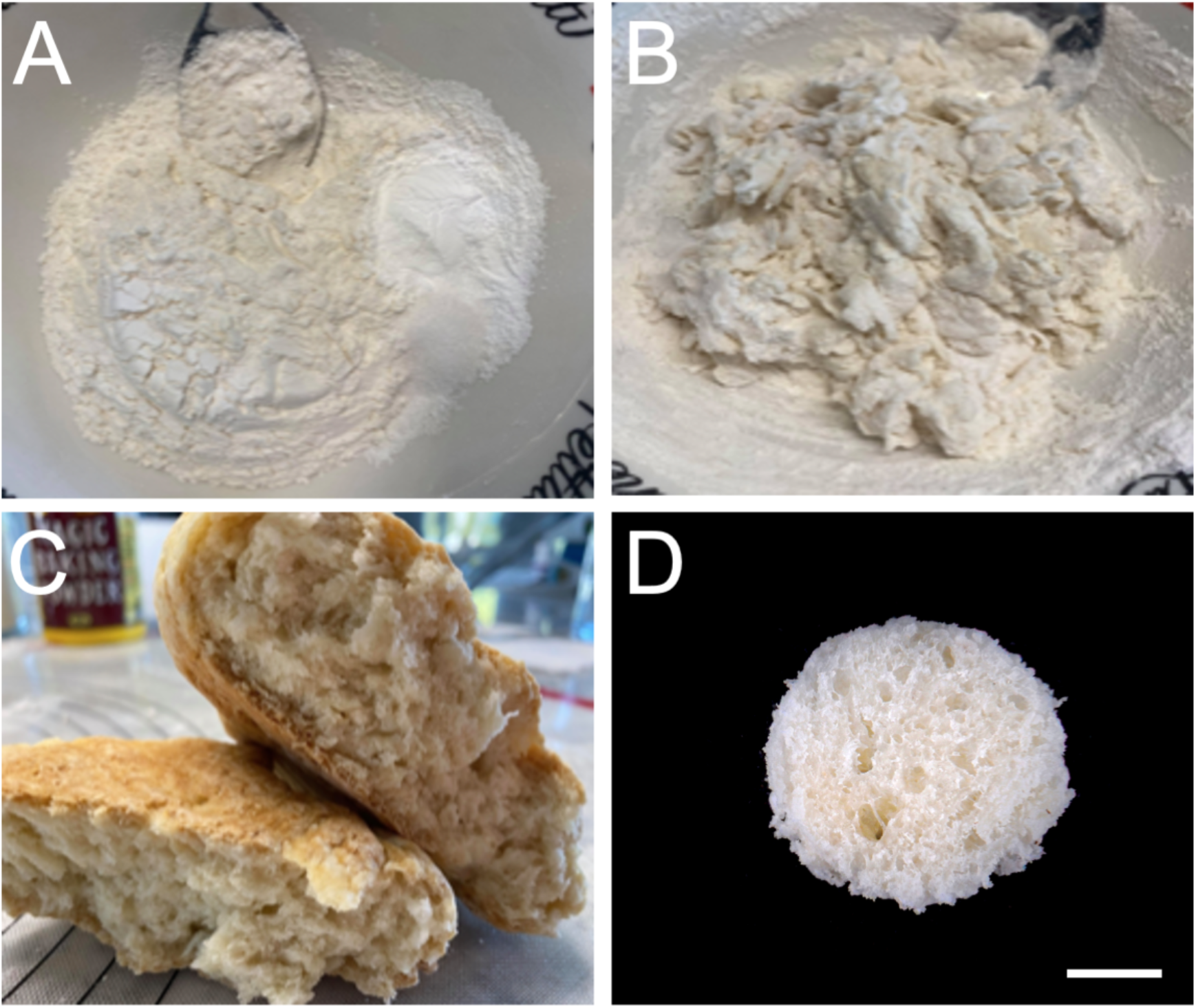
Fabrication of baked bread (BB) scaffolds. A) Dry ingredients (flour, salt, baking powder). B) Assembling the ingredients with warm water creates a dough. C) After baking the dough the bread crumb is utilized for the fabrication of scaffolds for cell culture testing. D) Utilizing a 6mm biopsy punch, 2.5mm thick cylindrical scaffolds can be created from the crumb (scale bar = 2mm). The porous nature of the scaffolds is considered to be sufficient to support mammalian cell culture.

### 3.2 Mechanical and structural stability of bread-derived scaffolds over time in culture conditions

We designed this study to assess cell proliferation and infiltration over the course of two weeks in culture. This would require the baked bread (BB) scaffolds to be continuously and completely submerged in cell culture media at 37°C for the entire length of time. We were concerned that the native structure of the scaffold may begin to soften significantly and/or decompose over this time course. Therefore, we also created scaffolds that were glutaraldehyde crosslinked (xBB) to create a more stable structure. The mechanical properties of BB and xBB scaffolds were then measured after initially submerging in cell culture media (Day 1), 24 hr (Day 2) and 288 hr (Day 13) in culture media at 37°C with no mammalian cells (Fig. 2A, B) (Day 1, 2 and 13 n-values were as follows, n = 20, 12, 19 for BB and n = 12, 11, 15 for xBB). The results demonstrate that initially, there is no statistically significant difference in Young’s modulus between the BB and xBB scaffolds due to the large variability (22.8 ± 9.3 kPa and 30.8 ± 9.9 p = 0.06716). However, there is a clear difference in mechanical properties of the BB and xBB scaffolds as a function of time. By Day 13, BB scaffolds are observed to soften significantly to 8.8 ± 3.8 kPa (p = 4.16854 × 10^−6^) compared to their state on Day 1. In contrast to Day 1, the xBB scaffolds do not soften significantly by Day 13, maintaining a value of 24.2 ± 8.1 kPa (p = 0.27115). Although there is a slight downward trend in the xBB scaffolds, it is clearly not as significant as the BB scaffolds. Furthermore, by Day 13, the BB scaffolds are also softer than their xBB counterparts (p = 2.32518 × 10^−6^). Regardless, in both cases the BB and XBB scaffolds maintain their porous morphology and structure after immersion in cell culture media as evidenced by both SEM and confocal imaging (Fig. 2C–F). This is unsurprising as the crosslinking with glutaraldehyde is performed after initial baking and the crosslinking is only serving to stabilize the scaffold, rather than alter its microarchitecture. SEM and confocal imaging reveals that the pore sizes observed in the material can vary significantly over the range of micrometers to millimeters.

**Figure 2.**
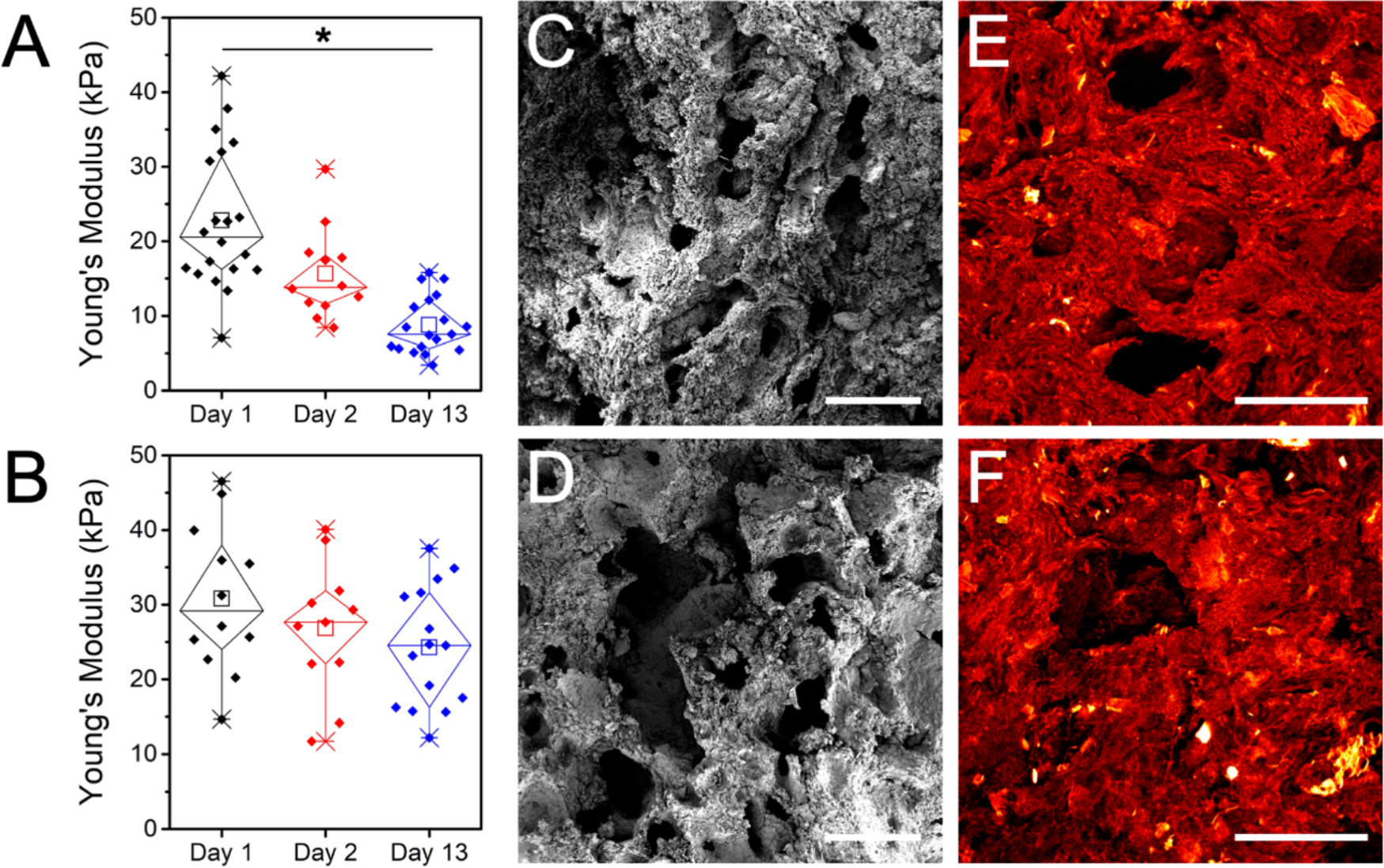
Preliminary analysis of the BB scaffolds after incubation in cell culture media. Mechanical analysis of A) BB and B) xBB scaffolds. Native BB scaffolds soften significantly by Day 13 compared to Day 1 (p = 4.16854 × 10^−6^). In contrast, the glutaraldehyde cross-linked xBB scaffolds do not exhibit any significant change in mechanical properties by Day 13 compared to Day 1 (p = 0.27115). The xBB scaffolds are also significantly stiffer than the BB scaffolds by Day 13 (p = 2.32518 × 10^−6^). SEM imaging at Day 13 of C) BB and D) xBB scaffolds reveals their porosity is maintained over time under culture conditions (scale bar = 1mm and applies to both). Maximum intensity Z-projections of confocal images acquired on congo red stained scaffolds also reveals the presence of pores in E) BB and F) xBB scaffolds (scale bar = 400um and applies to both).

However, admittedly glutaraldehyde is not an optimal cross-linker for food-based applications. As the scaffolds possess sufficient amounts of protein, well-known enzymes utilized in food processing, such as transglutaminase (TG) [26,27] (or other chemical crosslinkers and bread recipes), can be considered as alternatives. To validate the use of TG, we added it to the original formulation at a concentration of 1% (w/w) prior to baking. Once prepared the crosslinked scaffolds (tgBB) anecdotally felt more hydrated and robust. Mechanical testing confirms that the addition of TG significantly (p = 5.32719 × 10^−4^) increased the Young’s modulus of the scaffold in its dry form (161.7 ± 18.3 kPa, n = 12) compared to the dry BB scaffold (82.1 ± 7.1 kPa, n = 12) (Supplementary Fig. 1). Importantly, once in culture medium the tgBB scaffolds (n = 12) did soften as expected but reached a stiffness of 25.6 ± 4.3 kPa, consistent with the mechanical performance of the xBB scaffolds over time (Supplementary Fig. 1). Importantly, the mechanical properties of the tgBB scaffolds did not differ statistically from the xBB scaffolds (Fig. 2B) throughout their time course (p>0.8).

### 3.3 Porosity and pore interconnectivity

To further characterize the BB and xBB scaffolds, 1.3 × 1.3 mm confocal volumes were further analyzed to quantify the porosity of the scaffolds. Depth coded confocal image stacks of the BB scaffold reveals a highly complex structure with a number shallow pits and large pores which extend through the entire imaging volume (Fig. 3A). The pore structure of the BB and xBB scaffolds was largely composed of individual isolated pores and surface pits, as well as networks of interconnected pores underneath the outer surface (Fig. 3B). Confocal optical sections in a representative BB scaffold acquired 50um below the outer surface reveal the presence of both relatively flat continuous surfaces, as well as cross sections through individual pores (Fig. 3B). When imaging 150um below the surface, several important features are revealed (Fig. 3B). Firstly, there is clear evidence that underneath surface features on the exterior portion of the scaffold evidence of open volumes. Secondly, the bottom of open pores can be observed indicating that these are in fact pits which are isolated from the rest of the interior of the scaffold. Thirdly, it is possible to also observe how some pores open up into the voluminous space beneath the outer structure of the scaffold. A depth coded confocal image of an xBB scaffold is also shown for comparison (Fig. 3C). Importantly, pore sizes were observed to vary dramatically over the surface of the scaffolds. Representative images of BB (Fig. 3D) and xBB (Fig. 3E) are also presented which demonstrate the presence of very large (300-500um diameter) pores which can be routinely observed in these scaffolds. Although only a few data sets are discussed here, the data presented are representative of the scaffolds in general. However, it is important to appreciate that maximum intensity z-projections of confocal data tend to obscure the fact that the scaffolds possess both isolated pores as well as interconnected networks of pores. Likewise, it is important to keep in mind that SEM imaging is also a surface technique which may obscure deeper isolated and interconnected spaces within the scaffolds.

**Figure 3.**
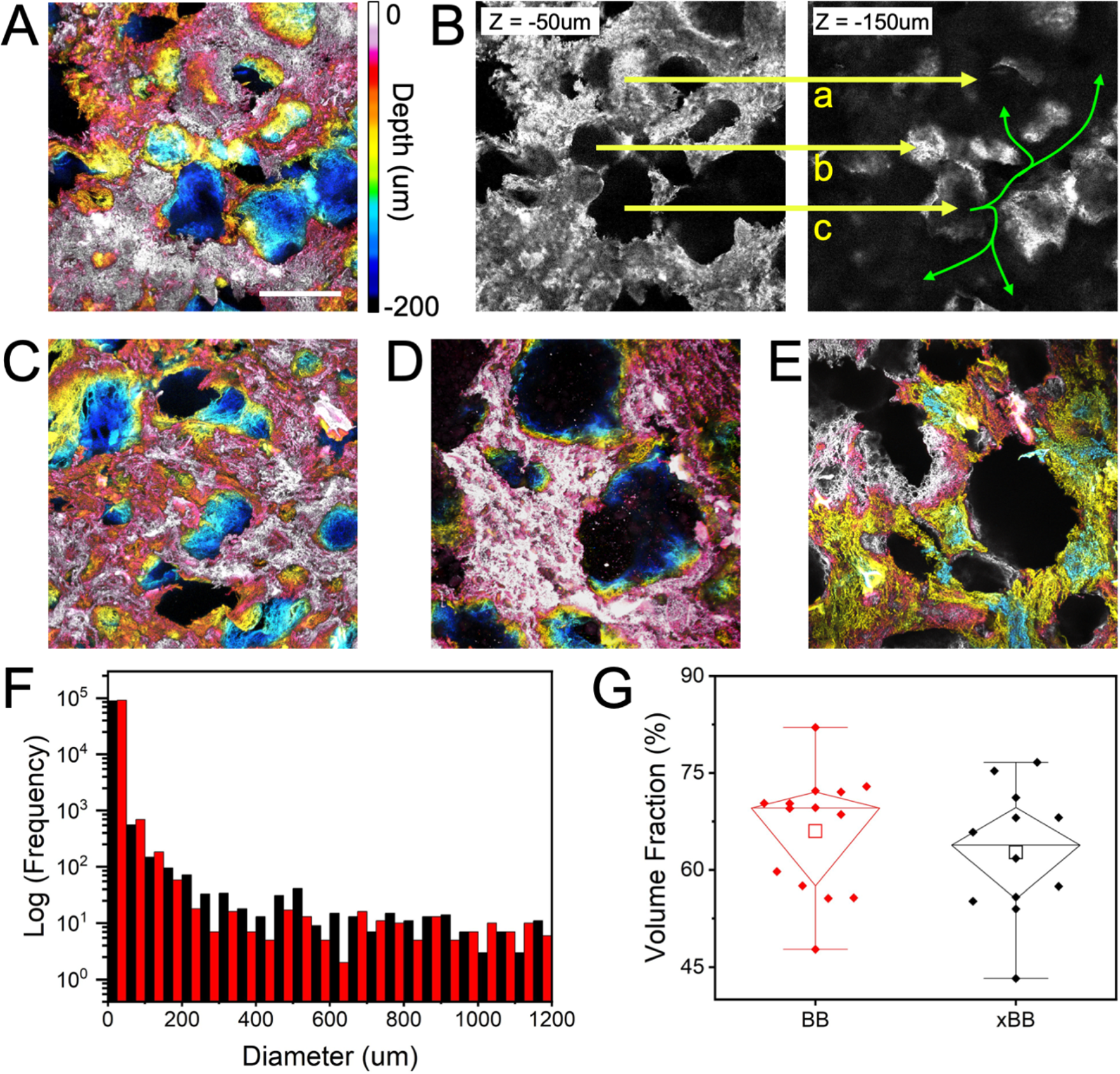
Characterization of BB and xBB scaffold microarchitectures. A) Confocal image of a BB scaffold where optical slices are coloured according to depth within the scaffold and maximum intensity Z-projected (color scale bar to the right of the image). The depth coded image reveals the complex porosity of the scaffold in more detail (scale bar = 300um and applies to all images, depth scale applies to all images). B) Single optical section of the data in (A) 50um (left image) and 150um (right image) below the surface. The left image reveals solid surfaces as well as open pores. A representative surface and two pores are identified by arrows a, b and c respectively (yellow). The arrows then point to the same region in the scaffold 150um below the surface (right image). Arrow “a” reveals how the solid surface in the left image covers a hollow void beneath (right image). Arrow “b” reveals the bottom surface (right image) of the pore identified in the left image. This particular pore is more accurately described as a pit and is isolated from the underlying open network. Finally, arrow “c” reveals how the bottom portion of a pore in the left image is actually open to an extensive interconnected volume of open space in the right image (green arrows). C) A representative depth coded maximum intensity Z-projection image of an xBB scaffold is presented for comparison to the BB scaffold in (A). Importantly, D) BB and E) xBB scaffolds also possess very large pore structures. F) A very large range of pore sizes can be extracted from confocal images of BB (red) and xBB (black) scaffolds. Notably, this distribution is likely incomplete as many pores can be macroscopic in nature and visible to the naked eye and such structures are not adequately captured in confocal data. G) A more useful measure of porosity is determining the volume fraction of empty space in the scaffolds. Both BB and xBB scaffolds possess statistically indistinct volume fractions of 66.0 ± 2.5 % and 62.7 ± 2.9 % respectively (p = 0.774565).

Standard image segmentation routines can be applied to the confocal data to isolate open volumes within the scaffold and quantify their size and volume fraction. Extracting the distribution of diameters from the data reveals a broad distribution of values ranging over three orders of magnitude (Fig. 3F). Importantly, the data does not follow a normal distribution and therefore estimates of mean and standard deviation are not particularly useful. We also note that as the pores of the scaffold are very often macroscopic and visible to the naked eye (Fig. 1D), the analysis of pore size is further complicated. At the cellular length scale (and high resolution of confocal microscopy) certain areas can appear quite flat but are in fact the surface of a larger pore within which are smaller pores. These large pores are not being adequately captured by the imaging area of the confocal microscope and the size distribution does not capture the full picture. Therefore, to gain some estimate of porosity we quantified the volume fraction of the scaffold occupied by free space (Fig. 3G). In this regard, confocal image volumes (n = 12-15 in each case) could be segmented and the fraction of the image volume occupied by the scaffold and free space can be easily quantified. BB and xBB scaffolds possessed a similar volume fraction of 66.0 ± 2.5 % and 62.7 ± 2.9 % (p = 0.774565). Interestingly, these volume fractions were similar to tgBB scaffolds (65.9 ± 2.4 %) (p > 0.8 in both cases).

### 3.4 Cell growth dynamics on BB and XBB scaffolds

Satisfied that the BB and xBB scaffolds were stable over time in cell culture conditions and media, we seeded them with NIH3T3 cells stably expressing GFP to assess cell proliferation. The scaffolds (n = 3 at each time point) were subsequently imaged using confocal microscopy at Day 2, 5, 7, 9, 11 and 13. The presence of cells after two days reveals that they adhere to both formulations and tend to initially invade inside of the scaffold pores (Fig. 4A, B). By Day 13 the cells have clearly proliferated forming a high density of cells (Fig. 4C–F). SEM imaging on the BB scaffold reveals cells covering the surface as well as proliferating inside of the pores (Fig. 4G). For each scaffold formulation (n = 3 per time point, per formulation), three randomly chosen 1.3 × 1.3 mm areas were imaged and the total number of cell nuclei were counted in each region and averaged (Fig. 4H). In both cases, the cell density on the scaffolds increases over time and eventually plateaus at about Day 9. The dynamics of cell growth do exhibit some variation during the experimental time course. Initially at Day 2, cell density on the BB and xBB scaffolds is the same (p = 0.9) at 548 ± 228 cells/mm^2^ vs 628 ± 428 cells/mm^2^, respectively. However at Day 5, the cell density on the BB scaffolds is significantly higher than the xBB scaffolds at 1682 ± 323 cells/mm^2^ vs 766 ± 260 cells/mm^2^, respectively (p = 8 × 10^−3^). While there is some variation in the proceeding time points, there are no statistically significant differences from Day 5 onwards. By Day 13 cell density on the BB and xBB scaffolds is the same at 2308 ± 339 cells/mm^2^ vs 1968 ± 494 cells/mm^2^, respectively (p = 0.94569). We also assessed the viability and metabolic activity of the cells at Day 1 and Day 13 with an alamar blue assay, which has been successfully used on 3D scaffolds [28] (Supplementary Fig. 2). The results are consistent with the cell count measurements presented here and provide more evidence that the scaffolds support cell growth over time.

**Figure 4.**
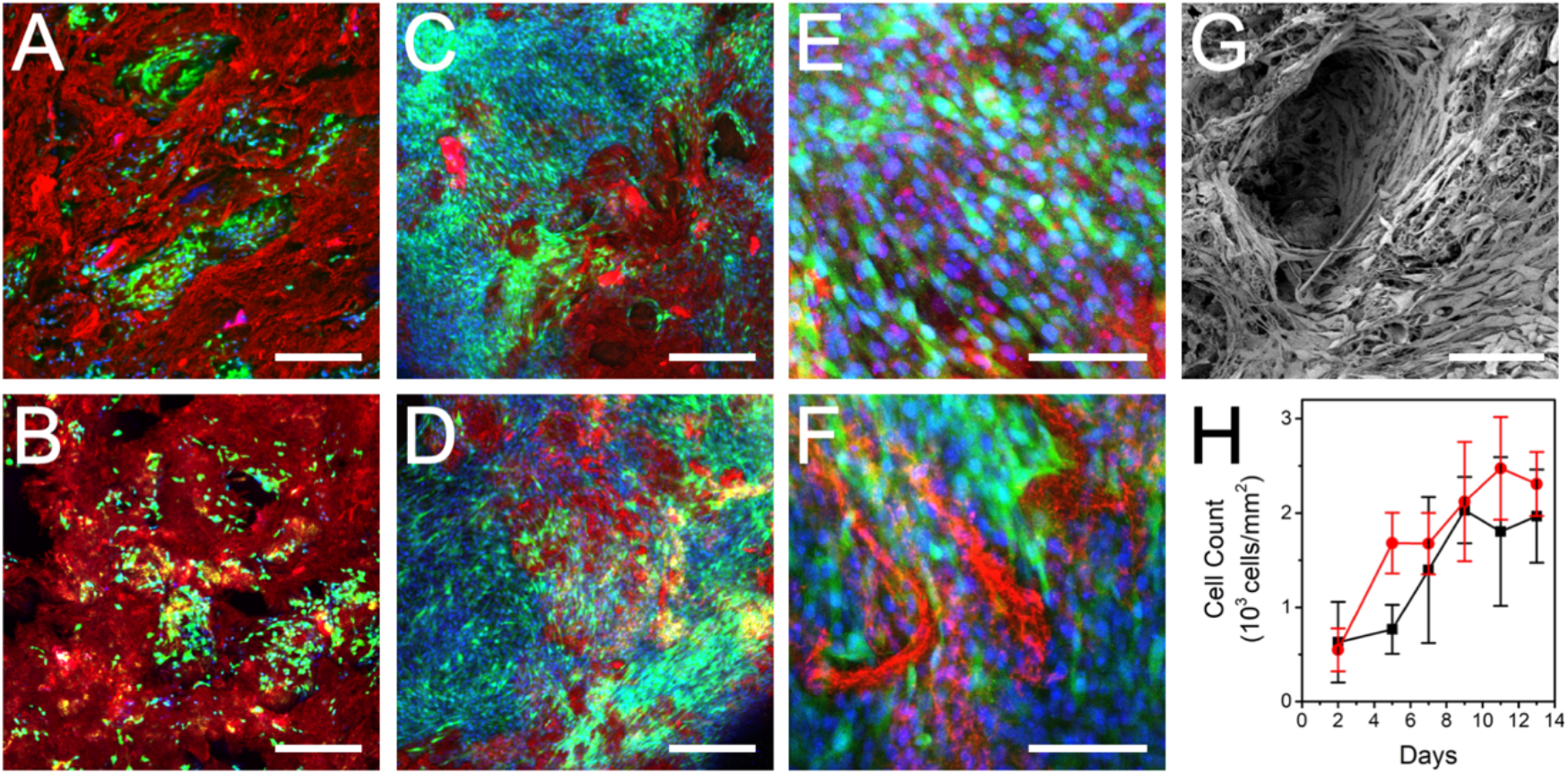
Microscopy and cell proliferation analysis of the BB and xBB scaffolds. On Day 2 after seeding the A) BB and B) xBB scaffolds reveal the presence of a low density of cells which are found on the surface and within cavities of each scaffold (scale bars = 300um) (blue = nuclei, green = GFP cells, red = scaffold). By Day 13 cell density has increased dramatically in C) BB and D) xBB scaffolds, with cells covering most of the available surface (scale bars = 300um). Higher magnification images of the E) BB and F) xBB scaffolds reveal the high density of cells (scale bars = 100um, all images are maximum intensity confocal Z-projections). G) SEM image of cells growing on a BB scaffold reveal how the cells are found inside of the pores of the scaffold as well as on free flat surfaces (scale bar = 100um). Similar results are also observed on xBB scaffolds (data not shown). H) Quantification of cell density on BB (red) and xBB (black) scaffolds reveals similar proliferation dynamics during the experimental time course. There is no significant difference in cell density between the scaffolds except at Day 5 (p = 0.008). This indicates that while the cells on the xBB scaffolds exhibit slower proliferation shortly after seeding, they eventually match the growth rate of cells on the BB scaffolds. By Day 13, the cell density on both scaffolds is not significantly different (p = 0.94569).

In order to assess the ability of the cells to penetrate into the scaffolds we cross sectioned them as described previously [3]. Briefly, on Day 13, a 6mm diameter, 2.5mm thick cylindrical scaffold was cut longitudinally with a microtome blade to produce two half cylinders. Subsequently, the cut side of the half cylinder could then be washed, fixed and prepared for imaging as described above. The results reveal that cells are indeed able to infiltrate into the deeper portions of the scaffold (Supplementary Fig. 3). However, as expected the cell density by Day 13 inside the scaffold is much lower than the outer portion [3,4,29]. This phenomenon is commonly observed in many 3D biomaterial scaffolds as it takes time to migrate deeply within a scaffold and due to diminished oxygen/nutrient diffusion the metabolic activity of these cells may be altered compared to cells closer to the exterior portion of the scaffold. As well, it is difficult to rule out how many cells were sheared off the scaffold surface during the sectioning process. Nevertheless, some cells were observed within the scaffold interior.

In addition to cell infiltration, we also stained scaffolds (n = 6) for the presence of the extracellular matrix protein fibronectin which is known to be produced by NIH3T3 cells [30]. In order to assess fibronectin deposition by the cells alone, we cultured them for one week to ensure scaffolds were not fully confluent. In this case fibronectin positive regions of the scaffolds tended to be localized with areas of higher cell density (Supplementary Fig. 4). Regions with very low cell density, or no cell density at all, did not demonstrate any significant positive signal for fibronectin. This result provides evidence that any fibronectin that might be present in the serum is not depositing on the scaffold. Alternatively, fibronectin from the serum may be depositing on the scaffolds but not at a concentration high enough to elicit a strong fluorescent signal.

### 3.5 BB and XBB scaffolds support the growth of multiple cell types

Finally, to perform an assessment of the generality of these scaffolds for multiple cell types we also cultured C2C12 muscle myoblasts (Fig. 5) and MC-3T3 pre-osteoblasts (Fig. 6). These cell types were both chosen as they are established model cell types commonly employed in research for tissue engineered scaffolds. In addition to cell growth, these cell types are also useful as they can be differentiated into muscle myotubes or osteoblasts which can serve as a useful tool for assessing their behaviour on a novel scaffolding material compared to other common scaffolding types [8,9,12,31].

**Figure 5.**
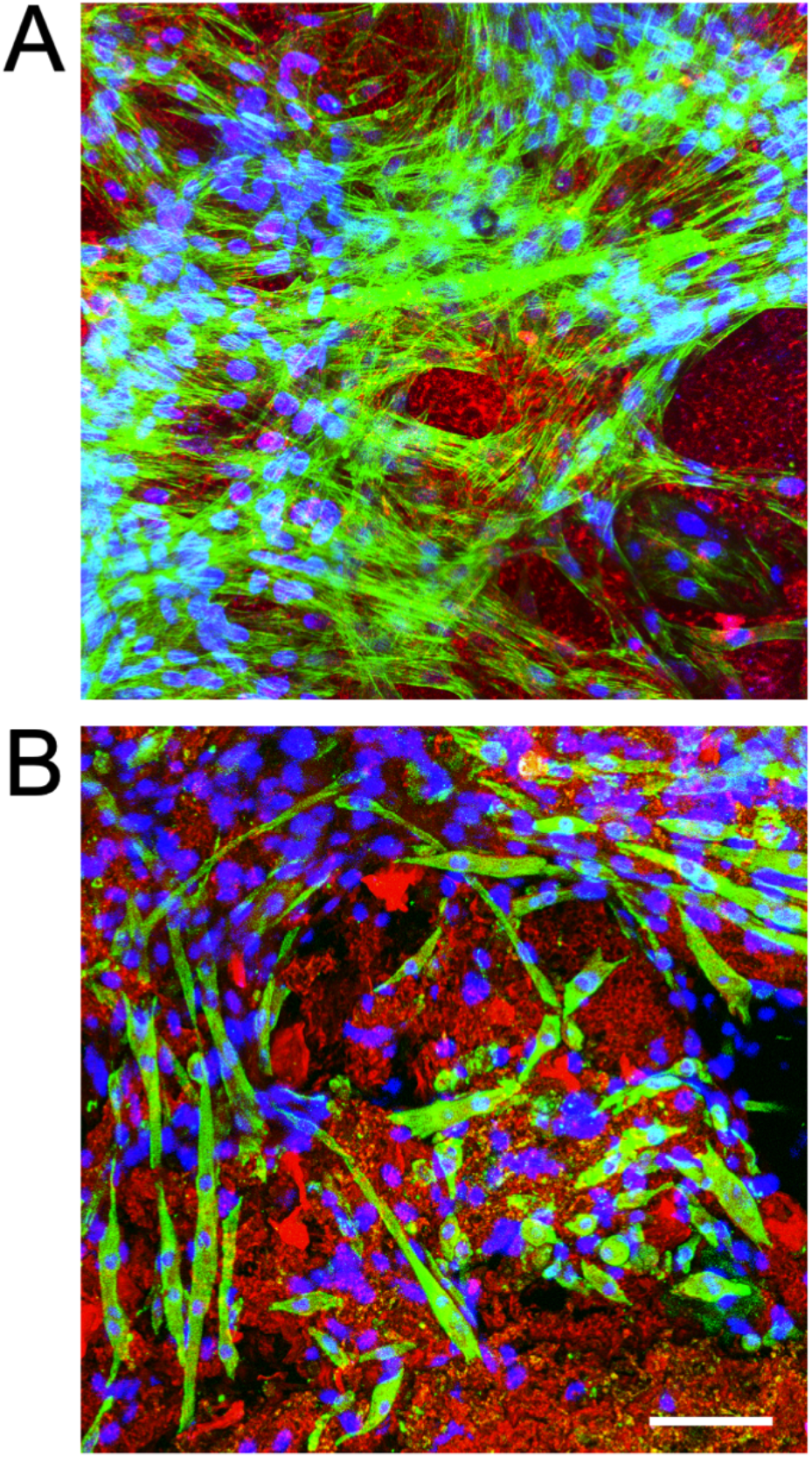
Culture and differentiation of C2C12 mouse myoblast cells on BB scaffolds. A) Mouse myoblasts after two weeks in culture (blue = nuclei, green = actin, red = scaffold). B) After differentiating the cells for two weeks they were observed to fuse into elongated multinucleated myotubes which appear green (blue = nuclei, green = myosin heavy chain, red = scaffold). Non-differentiated myoblasts are also visible as expected and appear as single blue nuclei which are not surrounded by any green staining. Scale bar = 100um and applies to both images, both images are confocal maximum intensity Z-projections.

**Figure 6.**
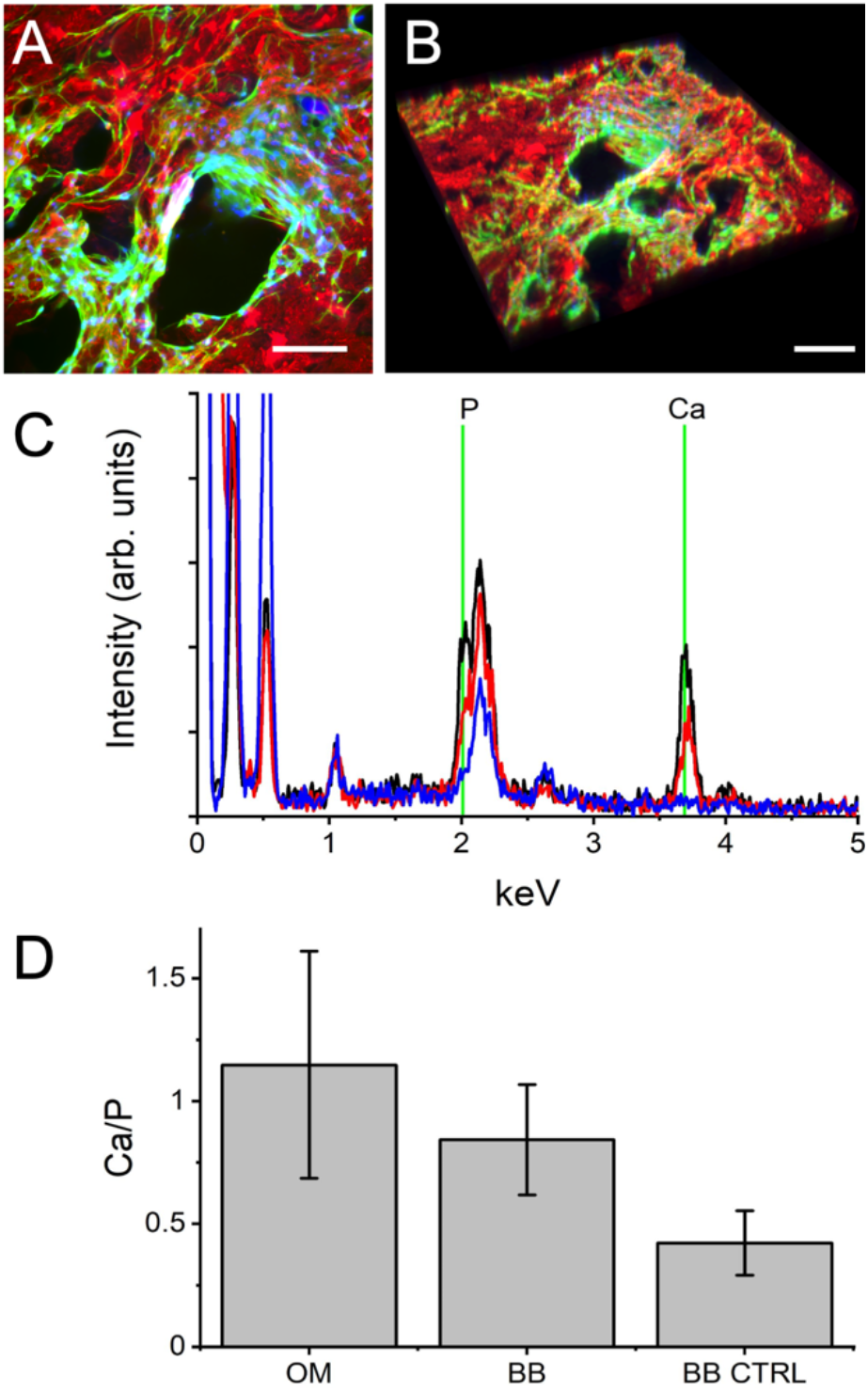
Culture and differentiation of MC-3T3 mouse pre-osteoblast cells on BB scaffolds. A) A confocal maximum intensity Z-projection of MC-3T3 mouse pre-osteoblasts after two weeks in culture on a BB scaffold, scale bar = 50um (blue = nuclei, green = actin, red = scaffold). B) A 3D reconstruction of a wider field of view of the data in (A) reveals the 3D nature of the scaffold, scale bar = 50um. C) Averaged EDS spectra of scaffolds containing cells cultured in OM (black), proliferation media (red) or a control scaffold with no cells (blue) (n = 3 in each case). A clear increase in P and Ca content in the samples observed from the control to proliferation media alone to OM. D) The Ca/P ratios reveal a clear increasing trend with differentiation, however due to variability this trend was not found to be statistically significant (p > 0.2 for all comparisons). The results are very important as they reveal how a given cell type may proliferate well on BB scaffolds but its capacity to differentiate appears to be impeded indicating that the scaffold may require further downstream optimization to support this particular cell type.

In the case of C2C12 cells, they were able to proliferate on both scaffold formulations in a manner consistent with the NIH3T3 cells. As they do not express GFP, we stained the actin cytoskeleton in addition to the scaffold and nuclei (Fig. 5A). C2C12 myoblasts were observed to migrate across the surface of the scaffolds and exhibit well defined actin stress fibres. These myoblasts are also a very common model for muscle myogenesis in which the cells proliferate to confluence after which they can be serum starved to stimulate their fusion and differentiation into multinucleated myotubes. This is a key early step in muscle tissue growth and formation. Here, we cultured the C2C12 cells on BB scaffolds (n = 12) for two weeks prior to switching them into differentiation media for an additional one to two weeks of culture. When differentiated myotubes were observed on the surface of the multi-well plates (some even spontaneously contracting) we prepared the scaffolds for staining. To identify differentiated myotubes we stained with an antibody against myosin heavy chain, a key indicator of differentiation [32,33]. Upon observation, myotubes were clearly observed on the scaffold surfaces consistent with more traditional substrates such as the standard plastic of tissue culture vessels (Fig. 5B).

In the case of the MC-3T3 pre-osteoblasts, the highly porous nature of the substrate is consistent with various other scaffolds encountered in bone tissue engineering [9]. In these cases, pre-osteoblasts are differentiated into osteoblasts which can mineralize porous 3D microenvironments [31,34]. Cells were first cultured for two weeks in proliferation media followed by switching into osteogenic media (OM) for an additional two weeks to differentiate. During differentiation with OM, or during biomaterial induced osteoinduction, this model cell line results in the formation of calcium and phosphorus rich mineral deposits on the underlying scaffold. In this case, after two weeks in proliferation media, cells were observed attached and proliferating on BB scaffolds in a manner consistent with the other cell types. As already noted, the scaffolds are highly porous and the cells are observed in the pores (Fig. 6A, B). After two weeks of proliferation, we switched the cells to OM for an additional two weeks. To assess the level of differentiation on the scaffolds, n = 3 scaffolds were prepared and energy-dispersive spectroscopy (EDS) was performed at three randomly chosen sites on each scaffold [9,34]. EDS spectra were acquired on differentiated cells, cells cultured for four weeks without OM and BB scaffolds alone (Fig. 6C). The EDS spectra clearly increased peaks occurring at 2.01 keV (phosphorus) and 3.69 keV (calcium) compared to the BB scaffold alone (Fig. 6C). Although the Ca/P ratio displays an increasing trend when comparing the BB scaffold alone and cells cultured with or without OM, the trend was not statistically significant (p > 0.2 in all comparisons) (Fig. 6D). Therefore, although some degree of mineralization is taking place in the presence of cells (with and without OM) the change is not significant enough to be confident that osteogenesis is occurring. This is an important result that indicates that while these bread-derived scaffolds do support cell proliferation, especially over a period of four weeks, there may be cell type specific issues that will require further scaffold optimization. Taken together, data presented here demonstrates that these scaffolds are likely to be compatible with a multitude of cell types. However, researchers should validate these scaffolds for each cell type that they intend to investigate as part of their own due diligence.

### 3.6 Metabolic activity of cells proliferating on BB and XBB scaffolds

To assess the relative health of all three cell types when cultured on the scaffolds in comparison to traditional 2D tissue culture plastic, we conducted both a lactate dehydrogenase (LDH) and glutathione (GSH) assay. The LDH assay is a common method for determining cytotoxicity and relies on the measurement of certain enzymes released by damaged or necrotic cells [35]. LDH is a cytoplasmic enzyme found in all cells and is released into the media when the plasma membrane is damaged or during cell death. Likewise, it has been well established that glutathione levels are a strong indicator of the ability of cells to detoxify and buffer against oxidative stress [36,37]. When total glutathione levels are observed to decrease this change can be measured to assess the degree of oxidative stress in a population of cells.

For the three cell types in question (NIH3T3, C2C12, MC-3T3) we examined both LDH and GSH in populations of cells cultured on BB scaffolds after two weeks in culture (Fig. 7A). In the case of LDH release, a statistically significant difference was observed on the 3D BB substrates compared to 2D substrates with comparable cell counts. For each cell type (n = 8 in each condition), a significant decrease in LDH levels of about 20% was observed for all three cell lines (p = 5.23 × 10^−3^, p = 2.94481 × 10^−6^, p = 1.27465 × 10^−6^ for the NIH3T3, C2C12 and MC-3T3 cells respectively). The results indicate that the cells on scaffolds are exhibiting less cytotoxicity, however, the observed levels on both tissue culture plastic and in the scaffolds are well within expected normal limits for many cell types.

**Figure 7.**
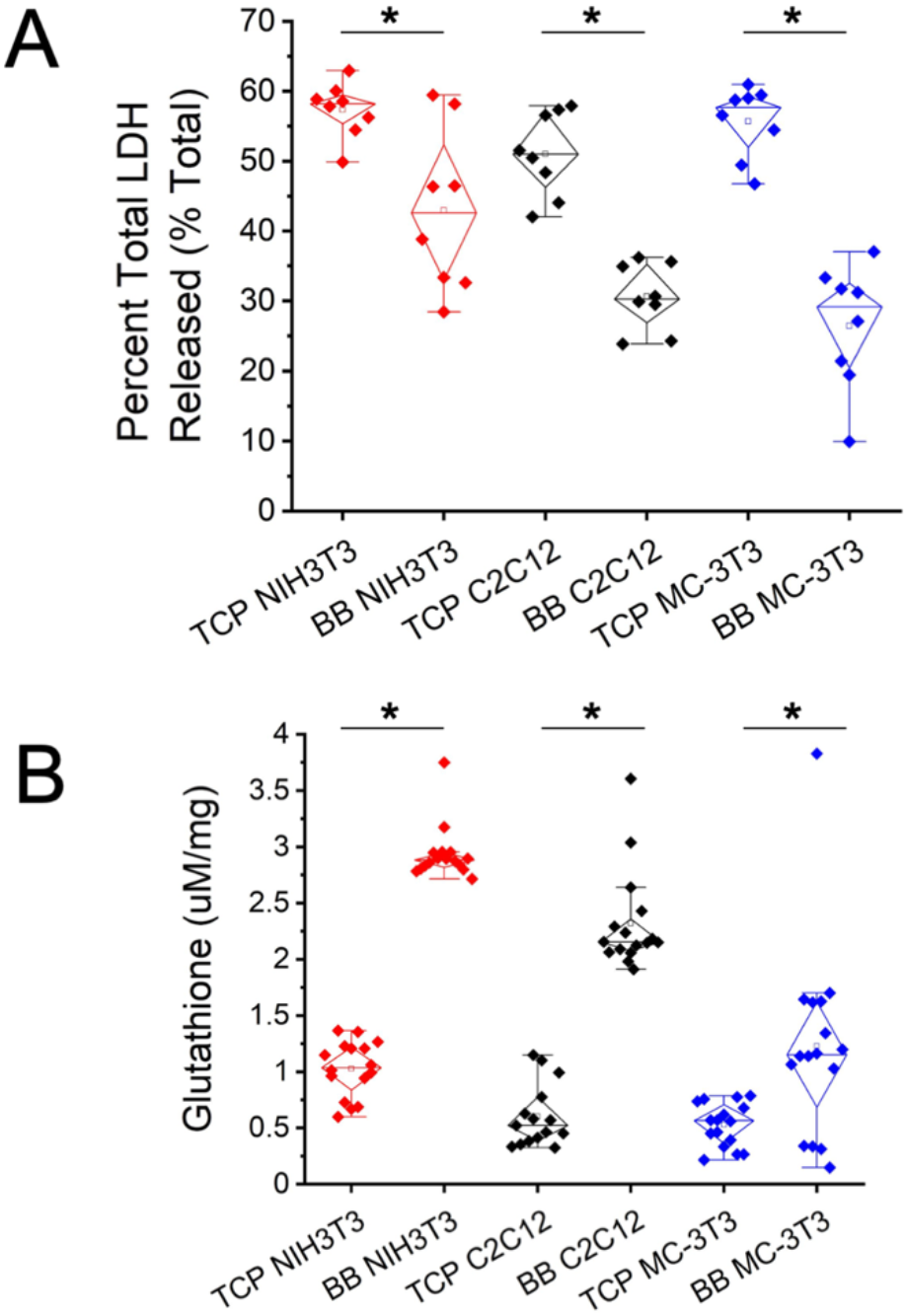
Lactate dehydrogenase (LDH) and glutathione (GSH) assays for assessment of cytotoxicity and oxidative stress when cells are cultured on tissue culture plastic (TCP) and BB scaffolds. A) LDH release during cell death is a well-known indicator of cytotoxicity. Importantly when cultured on BB scaffolds, cells actually exhibit a decreased levels of LDH (p = 5.23 × 10^−3^, p = 2.94481 × 10^−6^, p = 1.27465 × 10^−6^ for the NIH3T3, C2C12 and MC-3T3 cells respectively). B) Alternatively, total GSH is known to be reduced when cells are under oxidative stress. Consistent with the LDH data, cell populations cultured on BB scaffolds are observed to possess higher levels of GSH indicating lower levels of oxidative stress (p = 4.09462 × 10^−20^, p = 1.54964 × 10^−13^, p = 3.53 × 10^−3^ for NIH3T3, C2C12 and MC-3T3 cells respectively). While the trends are interesting, it is also important to note that the levels of LDH and GSH are well within normal and expected limits for healthy cells.

Consistent with these observations, the GSH assay suggested a similar conclusion. In this case we compared NIH3T3 (n = 16 in both conditions), C2C12 (n = 16 in both conditions), MC-3T3 (n = 15 and n = 16, on tissue culture plastic and BB scaffolds respectively). In all cases, total GSH was observed to increase by a factor of two to four when cultured on BB scaffolds as opposed to tissue culture plastic (p = 4.09462 × 10^−20^, p = 1.54964 × 10^−13^, p = 3.53 × 10^−3^ for NIH3T3, C2C12 and MC-3T3 cells respectively) (Fig. 7B). The increase in GSH is an indication that the cells are experiencing less oxidative stress when cultured on the BB scaffolds as opposed to the tissue culture plastic. While it may be possible that the BB scaffolds have sorptive properties it is unlikely that this will account for such significant differences. Furthermore, experimental measures were taken to ensure type I errors (i.e. false positive) were not committed during analysis by deprotienating samples with MPA and ensuring experimental controls were performed on unseeded BB scaffolds.

## 4. DISCUSSION

Bread is among one of the oldest foods in human history and its structure, mechanical properties and physicochemical characteristics have been studied extensively [38–41]. However, its applications for tissue engineering have not been explored to our knowledge. The porous architecture of the crumb has been used previously as a sacrificial material to produce scaffolds with potential applications for bone tissue engineering [42]. In this previous example, stale bread was immersed in a slurry of glass powder, which slowly impregnated the crumb. Once fully coated, high temperature thermal treatment removed any organic components, leaving behind a sintered silica-based porous scaffold. Although no direct supporting evidence was presented, the authors did speculate [42] that the mechanical and structural properties of such silica-based scaffolds might have relevance to applications in bone tissue engineering. In contrast, the use of proteins derived from wheat (for example gluten) have been explored extensively in a bioengineering context [13–15]. The viscoelastic properties of glutens, found in the majority of bread products, impart material properties that are of relevance to tissue engineering applications. As we have shown in our previous work on plant-derived biomaterials, novel scaffolds for tissue engineering applications can be created from unconventional source materials. Although the direct use of plant-derived scaffolds was unusual initially, their inertness and biocompatibility have now been explored by several groups [1–12]. Here, we explored the hypothesis that the porous nature of the crumb in bread might present another opportunity to develop a relatively simple scaffolding material which would support the growth of mammalian cells.

As we have demonstrated here, a modified Irish soda bread recipe can be utilized to culture cells on bread derived scaffolds for up to four weeks (Fig. 1). Although the data presented here have many implications for future and ongoing studies, here we attempt to keep our focus on demonstrating that these bread derived scaffolds possess reasonable mechanical properties, internal porosity to allow for 3D migration of cells and exhibit broad biocompatibility. Our first concern with bread-derived scaffolds was that they may soften significantly, or even decompose over time under standard culture conditions. While untreated BB scaffolds do soften (Fig. 2A) they ultimately stay within the range or normal tissue stiffnesses [43]. Moreover, we demonstrate that glutaraldehyde crosslinking (xBB) appears to stabilize the mechanical properties of the scaffolds so that they do not deteriorate over a two week period (Fig. 2B). The use of glutaraldehyde to reinforce the scaffolds (including a subsequent neutralization step) also did not appear to cause any dramatic cytotoxicity as the rate of cell proliferation was similar to the scaffolds that were untreated (Fig. 4H). However, Alamar blue viability assays (Supplemental Fig. 2) did detect a statistically significant drop in the number of viable cells on xBB versus BB scaffolds which was not consistent with image-based cell counting assays. Potential uncertainty in the quantification of cell density from microscopy data may arise due to limitations in imaging depth and penetration inherent in confocal approaches (Fig. 5C, D). As well, we suspect the porous nature of the scaffolds likely leads to the washing out of dead or non-adherent cells over the course of two weeks in culture as the media is changed daily [1]. Nonetheless, although these limitations arise with any 3D biomaterial there is still a significant amount of cell proliferation occurring on each scaffold over time, in three different cell lines.

When comparing the mechanical properties of the BB and xBB scaffolds (which generally vary from 10-30 kPa) with other traditional hydrogel scaffolds, the values tend to fall within the same range [29,43–45]. This leaves open the possibility of employing BB and xBB (or future variants) in a wide range of potential tissue engineering applications [29,45]. We also performed a pilot study of the use of a common food additive, transglutaminase (TG), and demonstrated that it has similar effects on the mechanical stability of the scaffolds (Supplemental Fig. 1). TG is an interesting enzyme as it is commonly in food preparation strategies for meat and baked goods. As it is food safe and edible, this presents a more appealing alternative to glutaraldehyde. However, to keep this study focused we did not delve into the numerous parameters (concentration, temperature, baking time, TG application protocols, etc) which can impact the performance and effects of TG on the stability, biocompatibility, structure and mechanical properties of the bread scaffolds. Future studies will focus on the systematic evaluation of TG in a variety of bread-derived scaffold preparations.

A detailed microscopic analysis of the scaffolds reveal they form highly porous 3D structures with an open volume fraction of about 65% (Fig. 2F/G) similar to a multitude of other scaffolding biomaterials developed previously [29]. Glutaraldehyde and TG crosslinking strategies did not have any appreciable impact on the volume fraction of the scaffolds. The sizes of the pores formed in the scaffold are widely distributed over three orders of magnitude, from micrometers to millimeters (Fig 2F). Importantly, we observed both isolated pits/pores as well as interconnected pores extending into the deeper portions of the scaffolds (Fig. 2). This results in surface pits/pores in which cells are essentially unable to migrate any deeper as well interconnected spaces in which the cells are able to migrate freely. Structural porosity in 3D scaffolds is an important feature for any biomaterial that might be utilized for tissue engineering applications. Porosity is created in our scaffolds by the action of sodium bicarbonate, the only leavening agent. Although not explored in detail here, the concentration of sodium bicarbonate and/or the use of other compounds may provide a route to further exert control over pore size which was highly variable in our scaffolds. Future efforts to resolve this variability might involve ball-milling and sieving the bicarbonate for discrete particle diameter ranges. Additionally, parameters of baking time, loading concentrations and other variables require further analysis and should be evaluated on an individual and systematic basis to further refine the control over the ultrastructure of these scaffolds.

After an extensive mechanical and structural analysis of the scaffolds we turned our attention to biocompatibility. In this study we demonstrate that multiple cell types (fibroblast, myoblast, pre-osteoblast) are all able to infiltrate and proliferate throughout the scaffolds over the course of two weeks (Figs. 4–6). NIH3T3 cells proliferate rapidly throughout the scaffolds and were even observed to deposit their fibronectin extracellular matrix (Supplemental Fig. 4). In addition, C2C12 myoblasts were also observed to proliferate, fuse and differentiate into myotubes over the course of a four week culture period. Multinucleated myotubes clearly expressing myosin heavy chain were observed on the scaffolds (Fig 5B). Conversely, MC-3T3 pre-osteoblasts were also able to be cultured on the scaffolds over a four week period. However, differentiation attempts did not provide clear evidence of mineralization of the scaffolds (Fig 6C/D). This important result highlights that while many cell types may be easily cultured on these scaffolds, there will likely be a need to optimize the scaffold formulation to support cell type specific functions. Although we decided to keep the current study focused on developing the primary proof-of-concept data to support the use of bread-derived scaffolds in cell culture applications, such parameters are being systematically studied in ongoing studies. Regardless of the ability to differentiate, evidence suggests that all three varieties of cells (NIH3T3, C2C12 and MC-3T3) responded favorably to the conditions for growth and proliferation within the BB and xBB scaffolds in contrast to 2D tissue culture plastic (Fig. 7). This may be ascribed to a marked reduction in anoxic conditions as cells within a 3D matrix are not in as close proximity to one another, however a variety of other factors are also possible. The sorptive properties of BB and xBB scaffolds merit further investigation for future studies; it is possible that reduction in ambient levels of nitrogenous waste products may greatly improve cell health in long term studies [46].

Here, the bread scaffolds were primarily composed of wheat-derived proteins, specifically gluten proteins, which likely impart stability to the scaffolds over time. The scaffold does not contain any animal-based products thereby reducing the cost and increasing the simplicity of the recipe albeit the scaffolds are cultured in complete media containing FBS. Though the purpose of this study is not to solve the longstanding issue of the use of animal-derived products (such as FBS) in cell culture based food applications, several serum substitutes are entering the market and have been reported in the literature which may eventually address this complex issue [47–49]. Importantly, our goal here is to establish the use of a new and unconventional scaffold which is intrinsically edible for potential future applications in cultured and plant-based foods. We do recognize that there are many challenges still to be met before it becomes widely accepted that cell cultured meats present a viable alternative to traditional animal agriculture methods [16,21,50,51]. One unaddressed challenge in this study is the potential for creating large full cut meat products [5,16,17]. It is unclear that the current formulation will support cell infiltration throughout centimeter scale scaffolds, especially as there is a heterogenous distribution of isolated and interconnected pore structures in the biomaterial. This of course is a challenge faced by all biomaterials regardless of their application in biomedical or food tissue engineering. We suspect that approaches currently being investigated by the field to potentially address this issue (microcarriers in suspension culture, layered assembly of thin scaffolds, composite biomaterials etc.) may offer a route to creating fully infiltrated scaffolds which may meet industrial manufacturing challenges [5,12,16,17,19].

Although the primary focus of the bread-derived scaffolds presented here has been on their potential use in future food applications, repairing and regenerating damaged or diseased human tissues by utilizing scaffolds to guide cell growth and differentiation is an area also under intense development [52,53]. Indeed, previously developed plant-derived scaffolds have applications in both human tissue engineering and food engineering applications [5–12]. Many remarkable scaffolds have been developed over decades with applications in soft tissue repair [54], neuroregeneration [55,56], bone tissue engineering [9,57], skin reconstruction [58,59], artificial corneas [60] and skeletal/cardiac muscle regeneration [8,61,62]. A number of approaches in these areas have led to the development of synthetic polymers or animal/insect proteins (collagen, fibronectin, silk, etc) with unique physical, chemical, electrical, optical and mechanical properties specific to each application. More recently, the science of tissue engineering and biomaterials is being applied to the generation of *in vitro* cultured meat [5,17,20]. This application has unique requirements as the final product must be food safe and edible, which creates additional considerations when developing new scaffolding technologies. Interestingly, the relatively simple bread-based biomaterials presented here may offer a route towards these goals. As the infrastructure to create bread at large scale is already present, there is the potential that this supply chain may fit within the broader manufacturing context of lab-grown meat. However, this is still highly speculative, as it will require deeper validation studies to fully determine if bread-derived scaffolds will be a viable solution for the cultured meat industry, especially at scale.

In this study, our goal was to demonstrate the possibility of utilizing highly available and accessible materials to create scaffolding capable of supporting mammalian cell growth *in vitro*. In a general sense, we demonstrate that it is possible to support the growth of multiple mammalian cell types using the methods presented here. The scaffolds support proliferation, some differentiation and do not elicit any substantive indications of cytotoxicity or oxidative stress. It should be recognized that while this proof-of-concept study validates the use of these bread-derived scaffolds in vitro, we caution the reader that it is highly likely different forms of scaffold optimization, including control over such aspects as porosity, microarchitecture, mechanical properties, and surface chemistry, etc. will be largely cell type dependent. Future studies by our group, and hopefully by other groups, will provide further systematic evaluation of all of these characteristics (and likely many more) in the context of specific applications.

## 5. CONCLUSIONS

We demonstrated the novel use of an ancient food product as a biomaterial for creating a 3D scaffold to support in vitro cell culture. A number of studies have demonstrated that plant proteins and polymers such as soy, zein, gluten, gliadin and cellulose etc [8,13,14] can be used in a variety of tissue engineering applications, though many involve advanced processing and purification methods. That said, the approaches developed here are highly complementary to previous studies on plant tissues and polymers. While traditional, synthetic biomaterials have often been preferred due to the control over their physicochemical properties, recent evidence has revealed new opportunities made possible with naturally derived biomaterials. Several types of cells were demonstrated to be both adherent and proliferate in long term culture. Although post-modification methods are typically required to control the architecture, chemical and mechanical properties of such biomaterials, the highly accessible nature and relatively low upfront material cost of bread-derived scaffolds make them an appealing choice for future cellular agriculture applications.

## AUTHORSHIP CONTRIBUTION STATEMENT

**Jessica T. Holmes:** Data curation; Formal analysis; Investigation; Methodology; Roles/Writing - original draft; Writing - review & editing. **Ziba Jaberansari:** Data curation; Investigation; Methodology; Roles/Writing - original draft; Writing - review & editing. **William Collins:** Data curation; Formal analysis; Investigation; Methodology; Roles/Writing - original draft; Writing - review & editing. **Maxime Leblanc Latour:** Data curation; Investigation; Methodology; Roles/Writing - original draft; Writing - review & editing. **Daniel J. Modulevsky:** Data curation; Investigation; Methodology; Roles/Writing - original draft; Writing - review & editing. **Andrew E. Pelling:** Conceptualization; Data curation; Formal analysis; Funding acquisition; Project administration; Resources; Supervision; Roles/Writing - original draft; Writing - review & editing.

## ACKNOWLEDGMENTS

This work was supported by grants to Andrew E. Pelling from the National Sciences and Engineering Research Council Discovery Grant, a Canada Research Chair, the Canada Foundation for Innovation and the Li Ka Shing Foundation. The authors would additionally like to give thanks for the support of Dr. Yun Liu of the Electron Microscope imaging core for her assistance in SEM imaging.

## DATA AVAILABILITY

The research data required to reproduce these findings is available upon request from the corresponding author.

