## Supplementary Material for "Homemade Bread: Repurposing an Ancient Technology for *in vitro* Tissue Engineering"

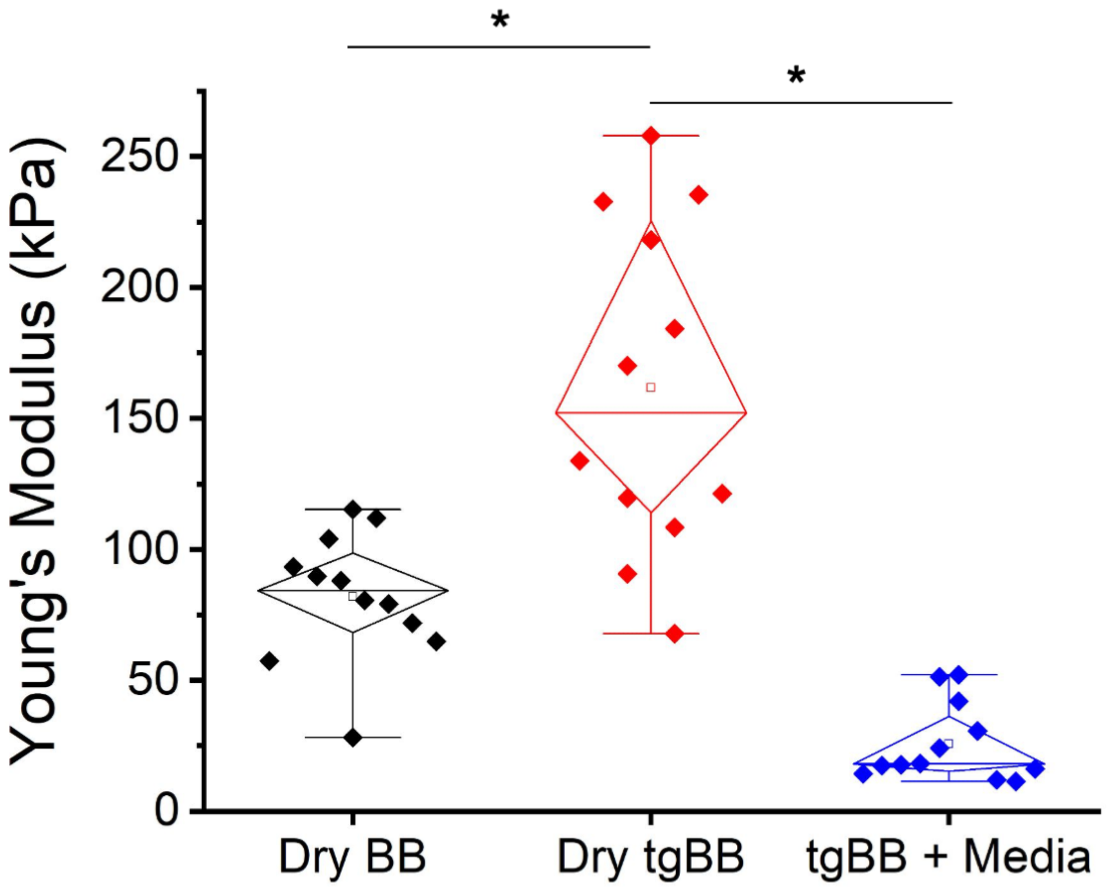


**Supplementary Fig. 1.** Young's modulus of dry BB scaffolds (black), dry TG crosslinked scaffolds (tgBB, red) and tgBB scaffolds after incubation in culture conditions (blue) measured on a CellScale Univert at 2%/sec to a maximum of 85% strain (the Young's modulus was determined from fitting the linear regime, typically 10-30% compression). The dry scaffolds are almost two times stiffer than the dry BB scaffolds ($p = 5.32719 \times 10^{-4}$). However, after being kept under culturing conditions the scaffolds softened in a manner consistent with the BB and xBB scaffolds. Importantly, over time the mechanical properties of the tgBB scaffolds in media were not statistically different from the xBB scaffolds ($p > 0.8$). The results support the potential use of TG as a crosslinker for potential applications in food processing which is likely preferable to the glutaraldehyde used in the xBB scaffolds.

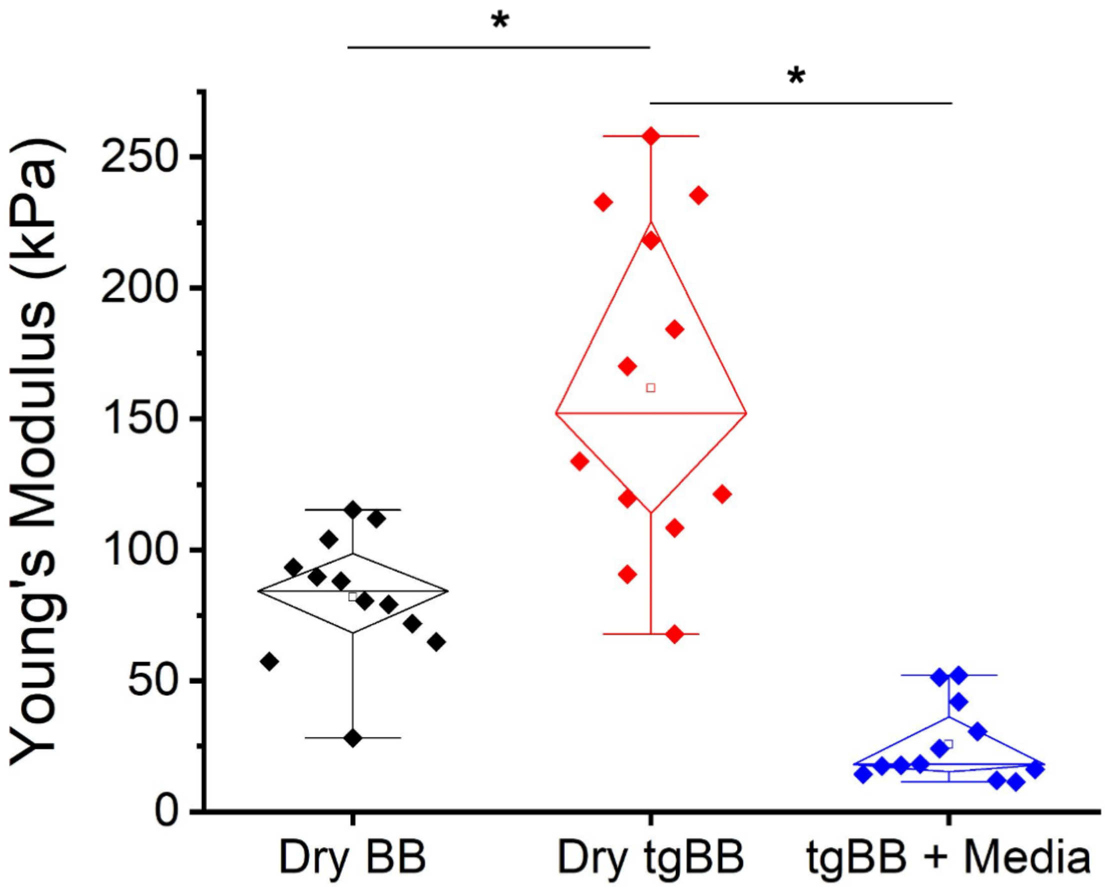


**Supplementary Fig. 2.** Alamar blue assay reveals the relative fold-increase metabolic activity of NIH3T3 fibroblasts after 1 and 13 days of culture on the BB (red) and xBB (black) scaffolds. For Day 1 and Day 13 BB scaffolds (n=32, n=27, respectively) cell viability increases significantly by a factor of ~4 on average ($p = 4.74779 \times 10^{-10}$). In contrast, on Day 1 and Day 13 xBB scaffolds (n=38, n=36, respectively) cell viability increases by a factor of ~3 on average ($p = 7.50372 \times 10^{-14}$). On both scaffold types the increase in cell viability is significant compared to the initial state. The BB scaffolds do display a large amount of variability in contrast to the xBB scaffolds, however on average there do appear to be more viable cells on these scaffolds ($p = 0.02143$). This may be due to the high degree of variability in the data, potential cytotoxic effects of glutaraldehyde or an unknown mechanism.

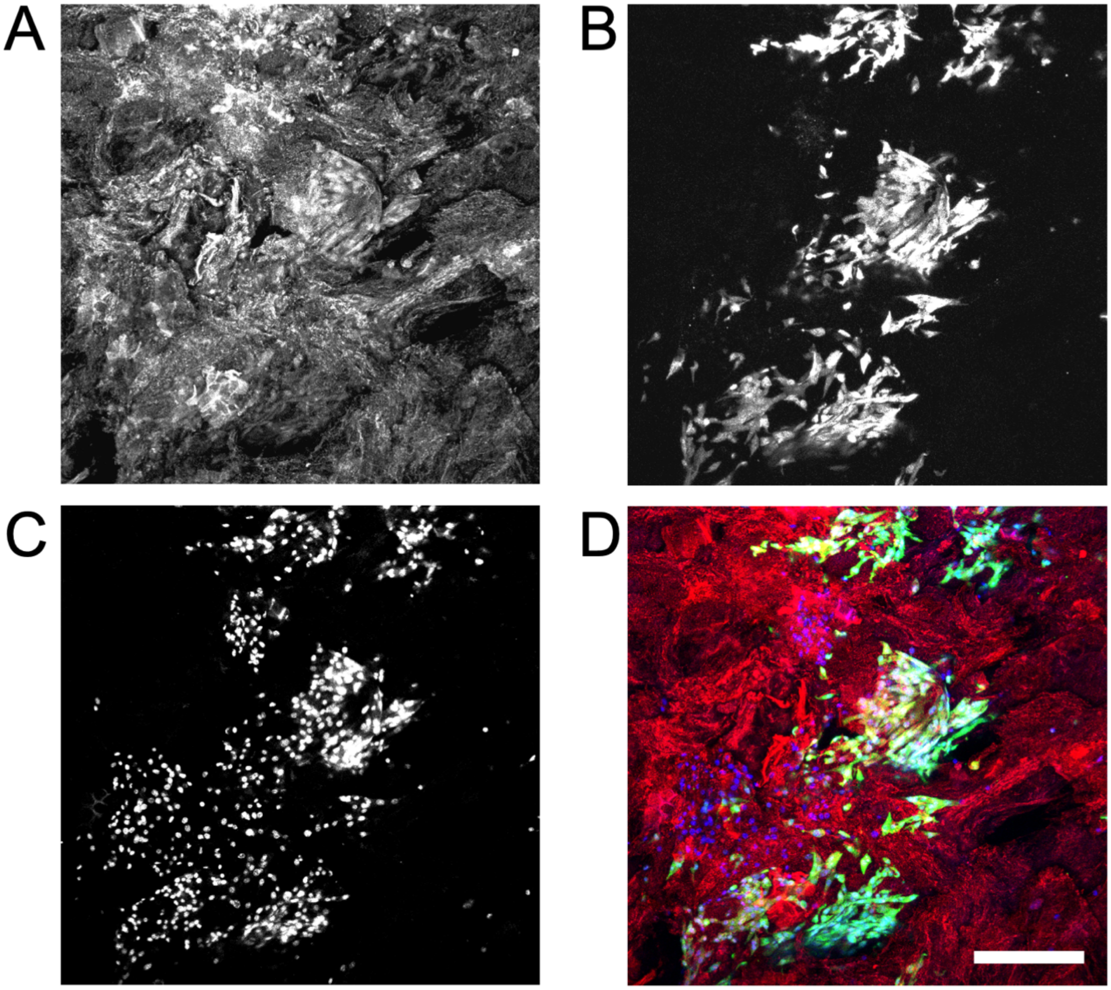


**Supplementary Fig. 3.** To investigate how deeply cells were able to penetrate into the BB scaffolds (n=3), they were cut in half on Day 13. A central region of the internal portion of the scaffold was then imaged. The A) scaffold; B) cell bodies; C) nuclei and D) merged image reveal that the cells are able to penetrate deeply inside of the scaffold (scale bar = 300um and applies to all). Consistent with other 3D biomaterials, cell density was clearly lower than the outer surfaces of the scaffolds when examined at the same point in the experimental time course (all images are maximum Z-projections of confocal data, blue = nuclei, green = GFP cells, red = scaffold).

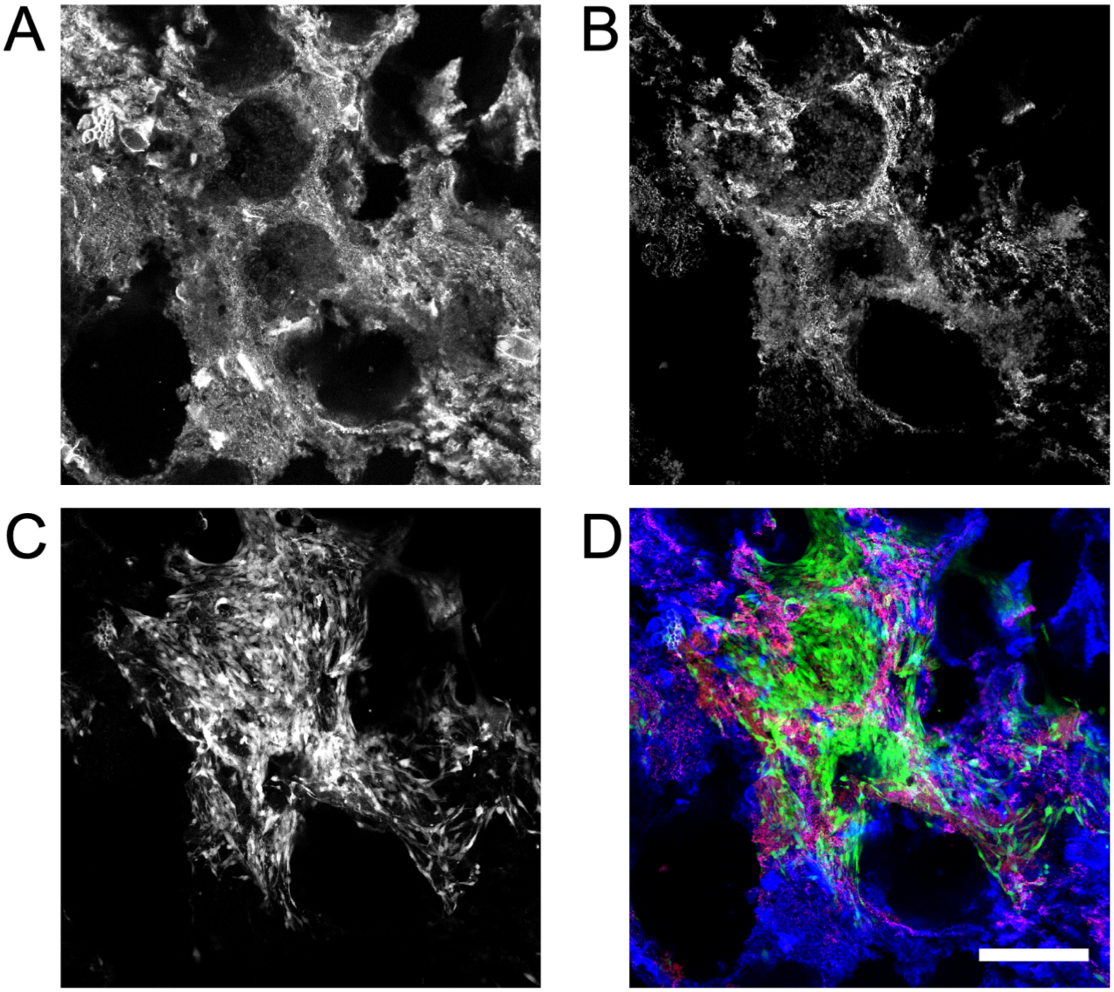


**Supplementary Fig. 4.** A) BB scaffolds containing B) NIH3T3-GFP cells after two weeks of culture were stained for C) fibronectin. D) A merged image showing the scaffold stained with calcofluor white (blue), NIH3T3 cells (green) and fibronectin deposits (red) (scale bar = 300um and applies to all). Notably, fibronectin deposition was not observed throughout the scaffold but tended to be largely localized to regions of significant cell density (all images are maximum Z-projections of confocal data, blue = scaffold, green = GFP cells, red = fibronectin).
